# Immunological memory to Common Cold Coronaviruses assessed longitudinally over a three-year period

**DOI:** 10.1101/2022.03.01.482548

**Authors:** Esther Dawen Yu, Tara M. Narowski, Eric Wang, Emily Garrigan, Jose Mateus, April Frazier, Daniela Weiskopf, Alba Grifoni, Lakshmanane Premkumar, Ricardo da Silva Antunes, Alessandro Sette

**Affiliations:** Center for Infectious Disease and Vaccine Research, La Jolla Institute for Immunology (LJI), La Jolla, CA 92037, USA; Department of Microbiology and Immunology, University of North Carolina School of Medicine, Chapel Hill, NC 27599-7290, USA; Department of Medicine, Division of Infectious Diseases and Global Public Health, University of California, San Diego (UCSD), La Jolla, CA 92037, USA

**Author notes:** These authors contributed equally (Ricardo da Silva Antunes, Alessandro Sette). Correspondence (R.d.S.A.), (A.S.).

## Abstract

Understanding immune memory to Common Cold Coronaviruses (CCCs) is relevant for assessing its potential impact on the outcomes of SARS-CoV-2 infection, and for the prospects of pan-corona vaccines development. We performed a longitudinal analysis, of pre-pandemic samples collected from 2016-2019. CD4+ T cells and antibody responses specific for CCC and to other respiratory viruses, and chronic or ubiquitous pathogens were assessed. CCC-specific memory CD4+ T cells were detected in most subjects, and their frequencies were comparable to those for other common antigens. Notably, responses to CCC and other antigens such as influenza and Tetanus Toxoid (TT) were sustained over time. CCC-specific CD4+ T cell responses were also associated with low numbers of HLA-DR+CD38+ cells and their magnitude did not correlate with yearly changes in the prevalence of CCC infections. Similarly, spike RBD-specific IgG responses for CCC were stable throughout the sampling period. Finally, high CD4+ T cell reactivity to CCC, but not antibody responses, was associated with high pre-existing SARS-CoV-2 immunity. Overall, these results suggest that the steady and sustained CCC responses observed in the study cohort are likely due to a relatively stable pool of CCC-specific memory CD4+ T cells instead of fast decaying responses and frequent reinfections.

## Introduction

Common Cold Coronaviruses (CCCs) are seasonal viruses comprising of two subtypes, namely α-coronaviruses (HCoV-229E and HCoV-NL63) and β-coronaviruses (HCoV-OC43 and HCoV-HKU1), that most frequently cause mild illnesses in humans ^1–5^. CCC are endemic viruses with widespread global distribution and have long circulated in humans. These CCC viruses are phylogenetically related to other coronaviruses that cause severe disease in humans, such as SARS-CoV-2, SARS-CoV and MERS-CoV ^6^.

CCC have been estimated to be responsible for up to 15-30% of pre-pandemic annual respiratory tract infections ^7–9^, with infections occurring most frequently in young children ^7,10,11^. CCC infections are associated with a clear seasonality, but infection can occur at any time of the year ^12–15^.

Whether immunity to CCC viruses is short or long lived has been debated with conflicting reports ^16–22^. Some discrepancies may be reconciled as some reports consider immunity as protection from re-infection while others consider protection from symptomatic disease. CCC infections are associated with generation of antibody titers widely detectable in the human population ^17,23–25^. However, little data is available regarding the frequency of memory T cell responses against CCC, and in particular their stability overtime. Understanding the steady state dynamics of CCC antibody and T cell responses in humans is of potential relevance in the context of the long-term evolution of the SARS-CoV-2 pandemic, in the context of the current scenario, where a large fraction of the human population is exposed and/or vaccinated.

Furthermore, it has been widely reported that CCC T cell responses are associated with some degree of cross-reactivity with SARS-CoV-2, and that this cross-reactivity can at least in part explain the pre-existing T cell memory reactivity recognizing SARS-CoV-2 sequences, observed in SARS-CoV-2 unexposed subjects ^25–29^. A putative role for CCC cross-reactive T cells in modulating COVID-19 vaccination and disease outcomes has been indicated by several independent studies ^26,30–33^. However, it is not clearly understood which factors in a given population, determine which individuals are associated with pre-existing SARS-CoV-2 T cell memory reactivity. Understanding the dynamics of CCC cross-reactivity with SARS-CoV-2 T cell responses is of potential relevance for understanding variations in COVID-19 disease severity, and also relevant to the potential development of pan-corona T cell vaccination ^34^.

Herein we performed a longitudinal analysis, over the course of 6 months up to 3 years, of CD4+ T cells and antibody responses to CCC and responses to other respiratory viruses, and chronic or ubiquitous pathogens. Overall, the results suggest that responses are readily detectable, and sustained over time.

## Results

### Frequency of CCC-specific memory CD4^+^ T cells are comparable to those for other common antigens

We studied PBMC samples from 32 participants of a *Bordetella pertussis* observational study ^35^. Three to seven longitudinal blood donations per donor, spanning time periods from 6 months to more than 3 years were available. All samples were collected in the 2016 to 2019 period (pre-pandemic). Subjects (9 male, 23 female) represented a range of ethnicities (14 Caucasian, 10 Hispanics, 7 Asian and 1 Black), with a median age of 24.5 years (range 18-35) (**Table 1**), and were recruited at LJI (La Jolla CA).

**Table 1.** Overall characteristics of the study cohort

| Number of Donors | Age (median and range) | Gender |  | Ethnicity |  |  |  |
| --- | --- | --- | --- | --- | --- | --- | --- |
|  |  | Male | Female | Caucasian | Hispanic / Latino | Asian | African American |
| 32 | 24.5 (18-35) | 9 | 23 | 14 | 10 | 7 | 1 |

CD4+ T cell responses to the four prototypical CCC viruses (NL63, 229E, HKU1 and OC43) were measured, using the Activation Induced Marker (AIM) and the OX40/4-1BB markers combination ^26^, which has been previously utilized to characterize viral responses and particularly SARS-CoV-2 CD4+ T cell responses ^36–40^. Responses to other respiratory viruses (influenza, RSV, and rhinovirus), chronically infectious viruses (EBV, CMV, and VZV), and ubiquitous bacterial vaccine antigens (TT and PT) were measured using specific peptide sets (**Table 2** and methods section). CD4^+^ T cell responses were measured in the 32 study subjects at the first-time point of the longitudinal series (**Fig. 1**). Significant antigen specific CD4+ T cell responses were detected for all four CCC epitope pools. Overall, 81.3%, 75.0%, 71.9%, and 78.1% of the donors were positive for NL63, 229E, HKU1 and OC43, respectively; median magnitude of CD4+ T cell responses was 0.089%, 0.083%, 0.078%, and 0.077% for NL63, 229E, HKU1 and OC43, respectively (**Fig. 1**). Similar reactivity across the 4 CCC were observed when considering SI responses ≥2 (**Fig. S1**) consistent with previous observations ^26^.

**Table 2.**
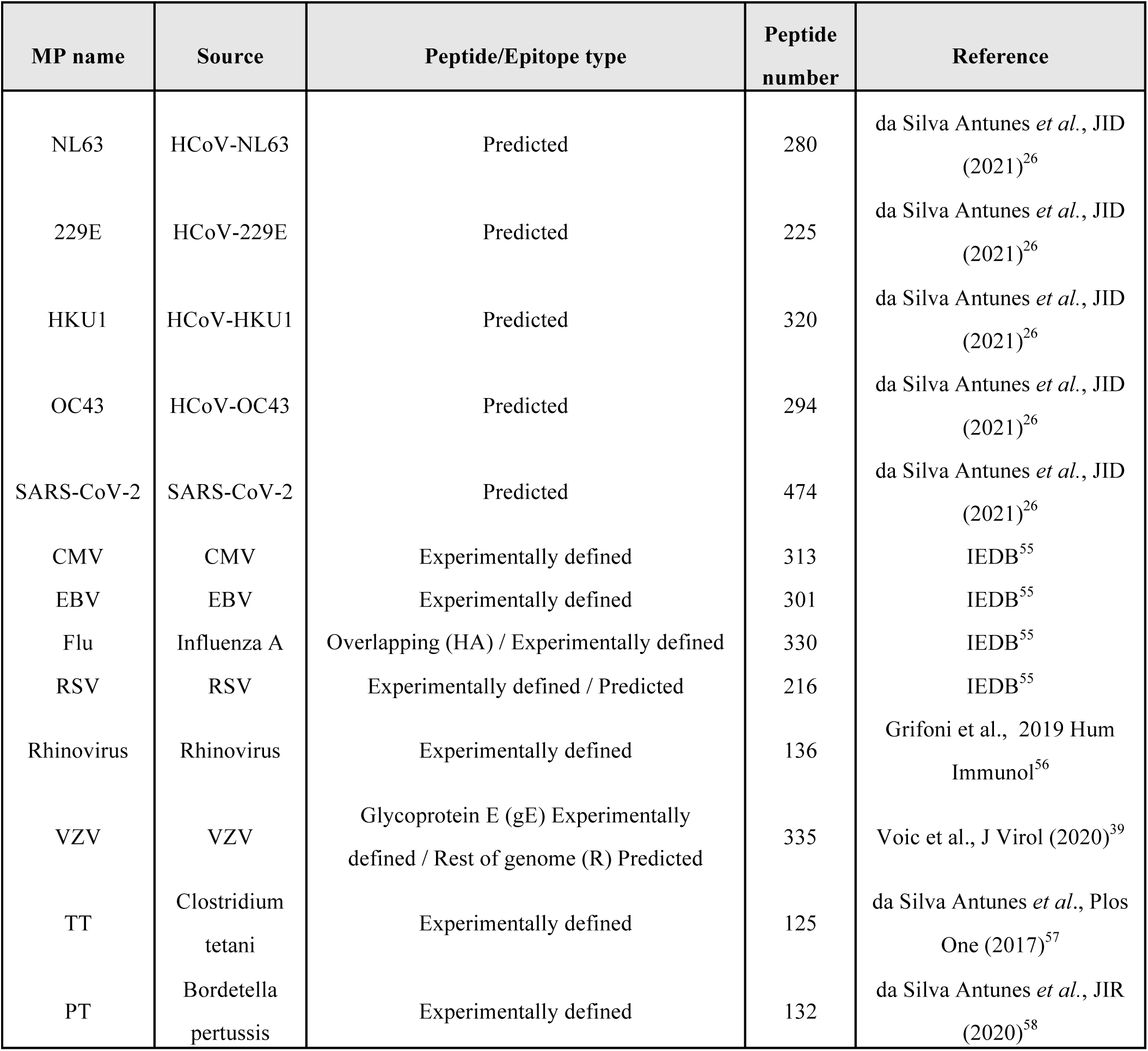
List of megapools (MP) used in this study

| MP name | Source | Peptide/Epitope type | Peptide number | Reference |
| --- | --- | --- | --- | --- |
| NL63 | HCoV-NL63 | Predicted | 280 | da Silva Antunes *et al.*, JID (2021)<sup>26</sup> |
| 229E | HCoV-229E | Predicted | 225 | da Silva Antunes *et al.*, JID (2021)<sup>26</sup> |
| HKU1 | HCoV-HKU1 | Predicted | 320 | da Silva Antunes *et al.*, JID (2021)<sup>26</sup> |
| OC43 | HCoV-OC43 | Predicted | 294 | da Silva Antunes *et al.*, JID (2021)<sup>26</sup> |
| SARS-CoV-2 | SARS-CoV-2 | Predicted | 474 | da Silva Antunes *et al.*, JID (2021)<sup>26</sup> |
| CMV | CMV | Experimentally defined | 313 | IEDB<sup>55</sup> |
| EBV | EBV | Experimentally defined | 301 | IEDB<sup>55</sup> |
| Flu | Influenza A | Overlapping (HA) / Experimentally defined | 330 | IEDB<sup>55</sup> |
| RSV | RSV | Experimentally defined / Predicted | 216 | IEDB<sup>55</sup> |
| Rhinovirus | Rhinovirus | Experimentally defined | 136 | Grifoni et al., 2019 Hum Immunol<sup>56</sup> |
| VZV | VZV | Glycoprotein E (gE) Experimentally defined / Rest of genome (R) Predicted | 335 | Voic et al., J Virol (2020)<sup>39</sup> |
| TT | Clostridium tetani | Experimentally defined | 125 | da Silva Antunes *et al.*, Plos One (2017)<sup>57</sup> |
| PT | Bordetella pertussis | Experimentally defined | 132 | da Silva Antunes *et al.*, JIR (2020)<sup>58</sup> |

**Fig. 1.**
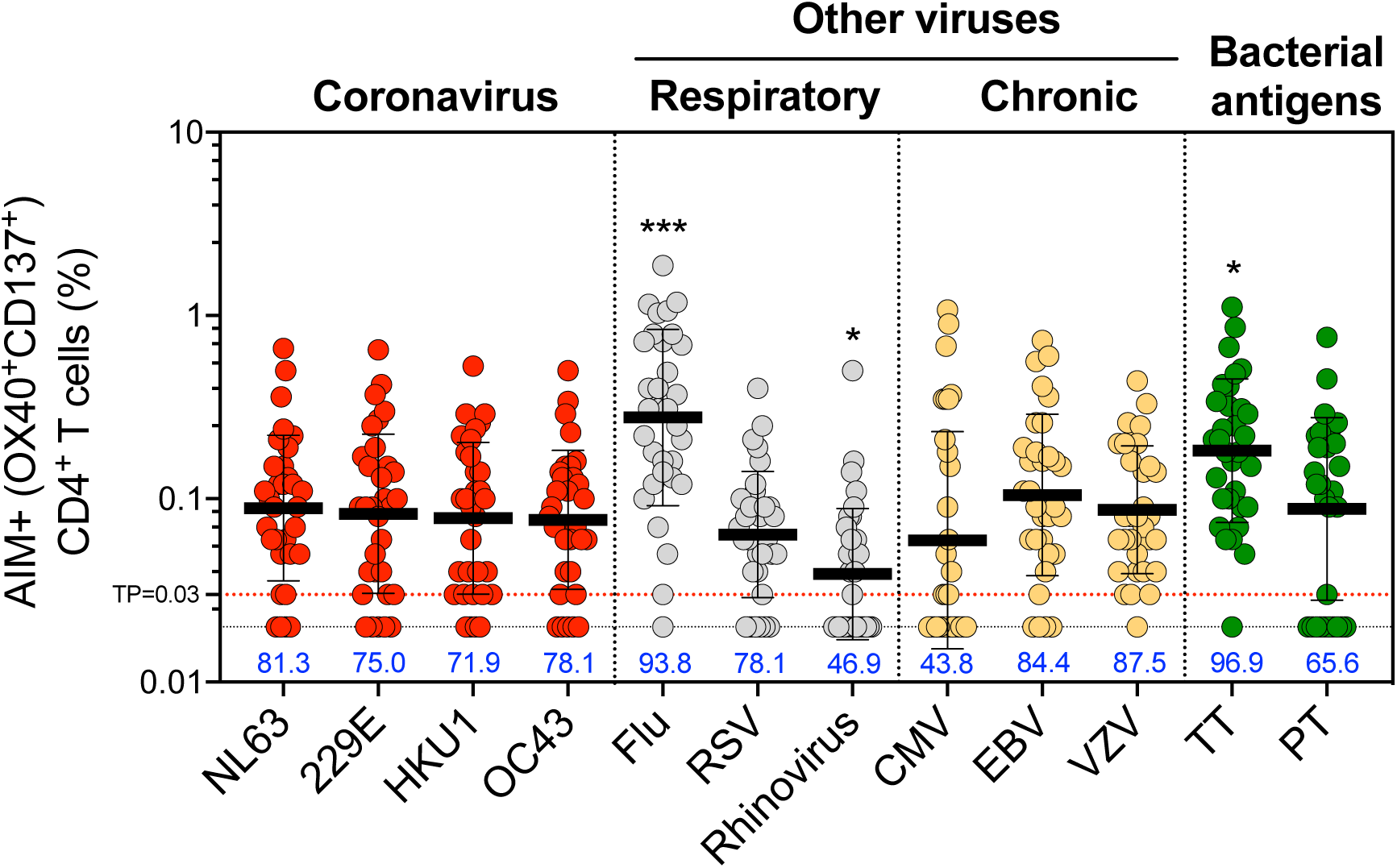
CD4+ T cell responses to four representative CCC are widely detectable in the study cohort and of similar magnitude to other pathogens. Common cold coronavirus (CCC) and several other human pathogens-specific T cell responses were measured as percentage of AIM+ (OX40+CD137+) CD4+ T cells after stimulation of PBMCs with peptides pools. Graphs show individual response of the four CCC (NL63, 229E, HKU1, and OC43) and other pathogens plotted as background subtracted against DMSO negative control. First time point of the longitudinal series is plotted (n = 32) and associated percentage of positive response for each antigen is indicated. TP=threshold of positivity. Data are represented as geometric mean and SD. Kruskal-Wallis test adjusted with Dunn’s test for multiple comparisons was performed and adjusted p values < 0.05 considered statistically significant. *, p < 0.05, ***, p < 0.001.

The CCC-specific CD4+ T cell reactivities were in the same range as those detected for the RSV, CMV, EBV, VZV and PT targets (**Fig. 1**). CCC-specific CD4+ T cell reactivities were 2-3-fold lower than Flu (p values ranging 0.0003-0.003 and p=0.01-0.04 for absolute and SI readouts, respectively) or TT (p values ranging 0.017-0.04 and p=0.003-0.004 for absolute and SI readouts, respectively) responses, and 2-fold higher compared to rhinovirus response (p values ranging 0.014-0.047 and p=0.024-0.036 for absolute and SI readouts, respectively) (**Fig. 1 and S1**).

As expected, CCC specific CD4+ T cells predominantly correspond to TCM and TEM memory compartments (defined as CD45RA-CCR7+ and CD45RA-CCR7-, respectively), with minimal contributions of naïve (CD45RA+CCR7+) or TEMRA (CD45RA+CCR7-) compartments **(Fig. 2)**. Similar phenotypes were associated with the AIM+ CD4+ T cells responding to the other antigen targets (data not shown). In summary, the data demonstrates that CD4+ T cell reactivity to 229E, NL63, HKU1 and OC43 was frequently detected in the study cohort, mediated by classic conventional memory cells and in the same order of magnitude as other viral antigens.

**Fig. 2.**
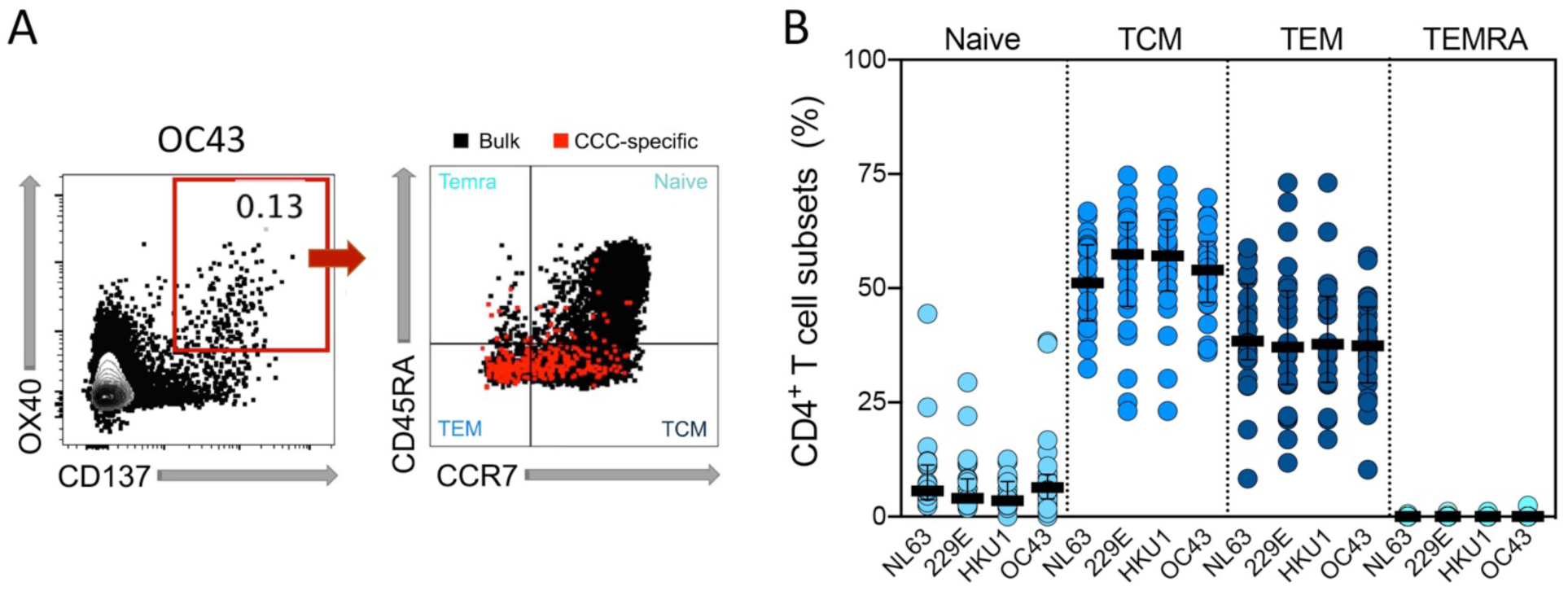
CCC-specific CD4+ T cells are largely classic memory CD4 T cells. CCC-specific CD4+ T cell subsets (Tn: CD45RA+ CCR7+, Temra: CD45RA+ CCR7-, Tcm: CD45RA-CCR7+ and Tem: CD45RA-CCR7-) were measured after stimulation of PBMCs with specific peptide pools. (**A**)Representative FACS plots, gated on the CCC-specific CD4+ T cells (red) measured as percentage of AIM+ (OX40+CD137+) from total CD4 T cells (Left), with the four subsets indicated in each quadrant for AIM+ cells (red) or total CD4+ T cells (black) (Right) are shown. (**B**)Percentages of T cell subsets from antigen-specific CD4+ T cells (OX40+CD137+) responding to the indicated pools of CCC and with SI>2 in each cohort (n = 32) at first time point are shown. Each dot represents the response of an individual subject to an individual pool and error bars represent median with interquantile range.

### Longitudinal analysis of CD4+ T cell reactivity to CCC and other antigens

We performed a longitudinal analysis of the levels of CD4+ T cell responses to CCC and other antigens. For each antigen, half-lives (t ½) were calculated based on linear mixed effects models using R package nlme ^41^, analyzing longitudinal responses for each individual. As shown in **Fig. 3A**, CCC CD4+ T cell responses were essentially steady over time (t ½ ranging to 244 years to no decline). Similarly, sustained responses were observed for the other antigenic targets **(Fig. 3B**). In particular, no decline was observed in the case of EBV, TT and PT or a modest decline (t ½= 7.4 years) in the case of influenza (**Fig. 3C**). Comparable patterns were observed for RSV, rhinovirus, CMV and VZV (**Fig. 3B**). Overall, these results indicate relatively constant and stable responses to CCC over the time considered, and in line to what observed for other antigens. To gain more insight in whether these apparently stable responses originate from frequent reinfections or long-lasting durable responses we determined the range of fluctuation of CD4+ T cell responses. This was done by first normalizing responses for each donor and each antigen, and then calculating the associated 5th-95th percentile range. We expected that responses to influenza, where yearly vaccination/exposures are relatively common, would fluctuate more than responses to other antigens, such as TT, for which natural exposure and re-vaccination is expected to be less frequent. The data in **Fig. 4A** indicates that this is indeed the case. Importantly, the patterns of fluctuation of CD4+ T cell responses, more specifically the 5^th^-95^th^ percentile range for each CCC viruses (ranging 0.31-3.85) were similar to that observed for TT (0.35-2.6), and lower than what was observed in the case of influenza (0.06-2.96). These data further indicated durable and constant CD4+ T cell responses to CCC overtime.

**Fig. 3.**
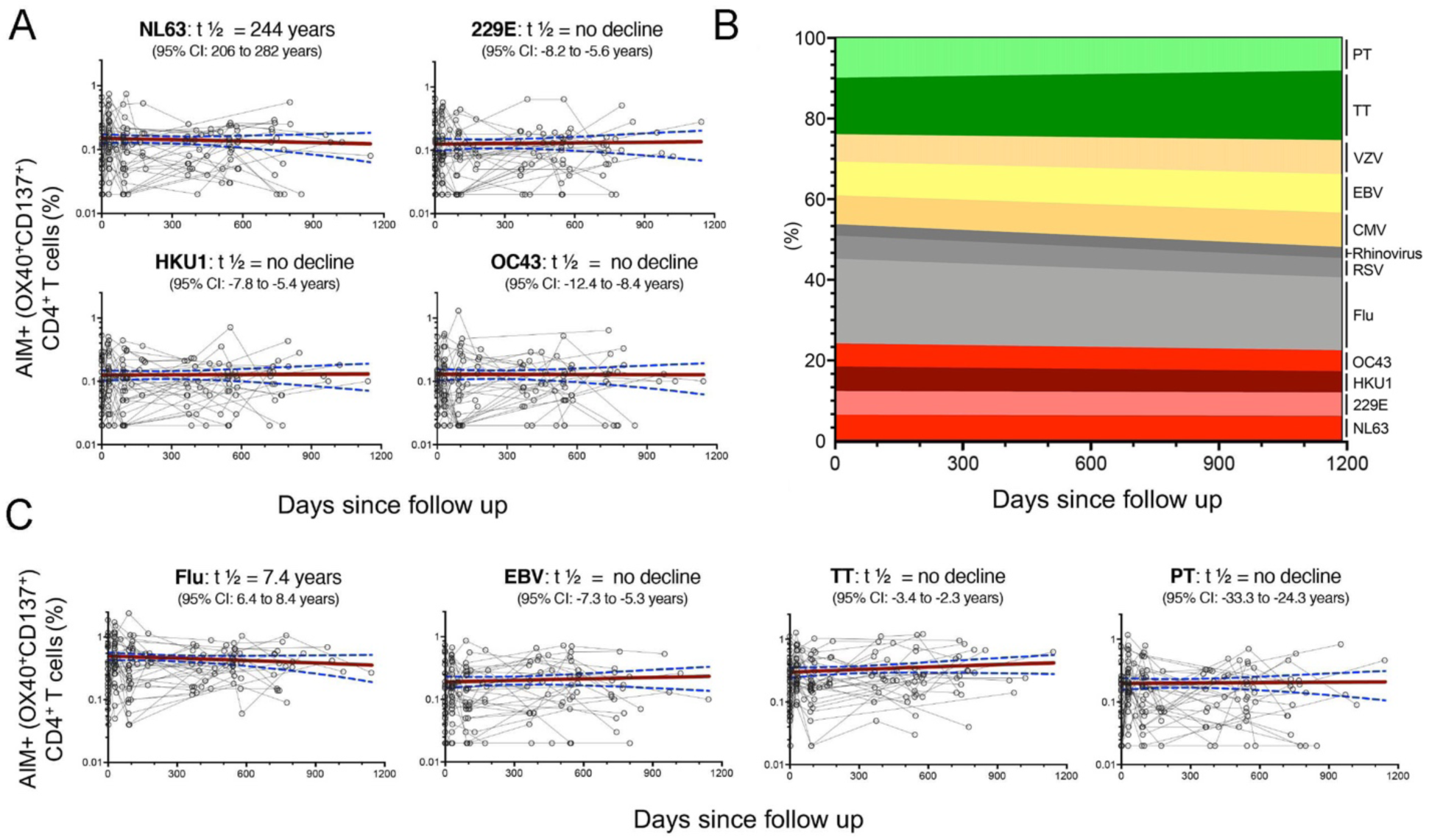
CD4+ T cells responses to CCC and other antigens are sustained over time. Antigen-specific T cell responses were measured as percentage of AIM+ (OX40+CD137+) CD4+ T cells after stimulation of PBMCs with peptides pools. Individual responses of the four CCC (**A**,**B**) or other pathogens (**B**,**C**) are shown. (**A**,**C**) Graphs show responses plotted with all time points of the longitudinal series connected with lines for each subject (n = 32). The red line represents the median fitted curve from a nonlinear mixed effects model of longitudinal responses among those with a positive response at ≥ 1 time point, with 95%CI shown in blue dotted lines. t ½ calculated based on linear mixed effects model using R package nlme ^41^, t^1/2^ is shown as the median half-life estimated from the median slope with the associated 95% CI indicated. (**B**) Longitudinal occurrence of each individual pathogen response distributed in overall percentage (sum of all absolute responses) in relation to the days since follow up.

**Fig. 4.**
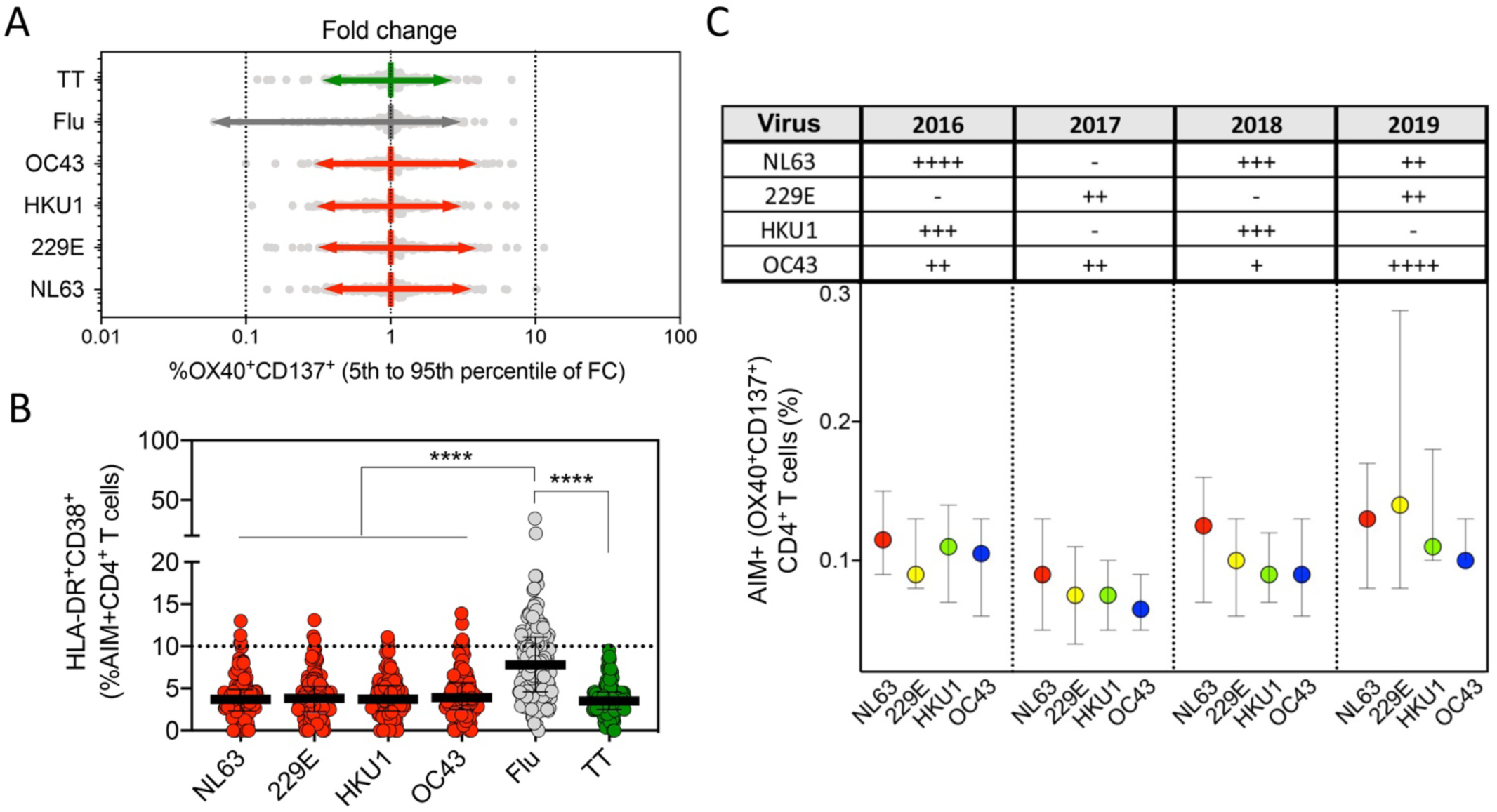
CCC-specific CD4+ T cell responses are stable, not associated with recent activation or yearly changes in prevalence of CCC infections. (**A**) The range of fluctuation of CD4+ T cell responses was determined by calculating the fold change of antigen-specific AIM+ (OX40+CD137+) CD4+ T cells. For each antigen, AIM+ CD4+ responses at every time point were normalized to median of total longitudinal responses for each donor (n = 32), and the 5^th^ to 95^th^ percentile range calculated. (**B**) Graph shows CCC, influenza and tetanus specific CD4+ T cell responses associated with recent activation measured by calculating the % of HLA-DR^+^CD38^+^ from AIM+ (OX40^+^CD137^+^) CD4+ T cells at all time points of the longitudinal cohort. Each dot represents the response of an individual subject (n=32) to an individual pool at a single time point. Median and interquartile range are represented. (**C**) The prevalences of CCC infections in the west and midwest regions during 2016-2019 were categorized according to the percent of positive rates from total tests performed ^12,14^: -, <1%; +, 1-2%, ++, 2-5%; +++, 5-8%; ++++, >8%, and results summarized in the table insert. CCC-specific CD4+ T cell responses for the four CCC were ploted as function of the yearly incidence (2016-2019) in the graph below. Median and interquartile range are represented. Kruskal-Wallis test adjusted with Dunn’s test for multiple comparisons was performed and adjusted p values < 0.05 considered statistically significant.

### HLA-DR^+^CD38+ expression and periodicity of CCC antigen-specific CD4+ T cells

The expression of the HLA-DR and CD38 markers is associated with recent *in vivo* activation ^26,42,43^. **Fig. 4B** indicate that CD4+ T cells responding to the CCC peptides are associated with a 3.7-3.9% range of HLA-DR^+^CD38^+^ AIM+ CD4+ T cells (95% confidence interval of 0.2-9.6%). Only few data points were above a threshold of high reactivity (>10%), which we previously associated with recent infection by SARS-CoV-2 of a cohort of COVID-19 convalescent subjects ^26^. Interestingly, influenza-specific CD4+ T cells were associated with a median of HLA-DR^+^CD38^+^ AIM+ CD4+ T cells of 7.8% (95% confidence interval of 2.2-16.0%), and 30% of the data points (donor/time point instances) were associated with values of 10% or higher. In the case of TT, the percent of HLA-DR^+^CD38^+^ AIM+ CD4+ T cells was similar to CCC (3.5% with a 95% confidence interval of 1.1-7.4%), with no data point above 10%. These data are consistent with relatively more frequent exposure to influenza as compared to TT responses, and are not consistent with frequenter re-exposure to CCC of the study cohort, within the time frame of the longitudinal study.

The data above are derived from longitudinal samples collected in the pre-pandemic 2016-2019 calendar years period. Epidemiological data is available regarding the circulation of CCC in those years, pertaining to the West and Midwest regions ^12,14^. If the CCC responses detected were short-lived responses, resulting from frequent re-exposure, we expected that the responses would mirror the CCC circulation pattern when segregated by year. When CCC CD4+ T cell responses were plotted as a function of the year in which the blood donation was obtained, no significant associations using a multiple comparison test, were observed with the yearly incidence of each individual CCC over the same period or when comparing different CCC responses within a year (**Fig. 4C**). Overall, these data suggest that CD4+ T CCC-specific responses are not associated with recent activation or frequent yearly reinfections.

### CCC circulating antibodies over time confirmed durable antibody titers and infrequent reinfections

Matched plasma samples were tested for binding of immunoglobulin (Ig) to recombinant Spike RBD antigens from CCC as previously reported ^44^. More specifically, we measured the RBD IgG levels by the area under the curve (AUC) in titration experiments for each individual sample (**Fig. S2)**.

All subjects were IgG seropositive for the four CCC (NL63, 229E, HKU1 and OC43) at the first and subsequent blood donations, consistent with their high prevalence in human adult populations ^12,17,45^ (**Fig. 5A**). Durability assessments of circulating antibody titers were performed as based curve fits to model ^41^, similarly to the assessments of CD4+ T cells responses above. CCC titers were sustained overtime for all the 4 CCC (**Fig. 1B**); NL63 and 229E titers showed no decline throughout the study, while a modest decline was observed for HKU1 and OC43 with a t_1/2_ of 13.1 and 8.6 years, respectively. The stability of CCC-specific IgG responses indicates that the cohort analyzed was not associated with frequent reinfections during the time considered in the study.

**Fig. 5.**
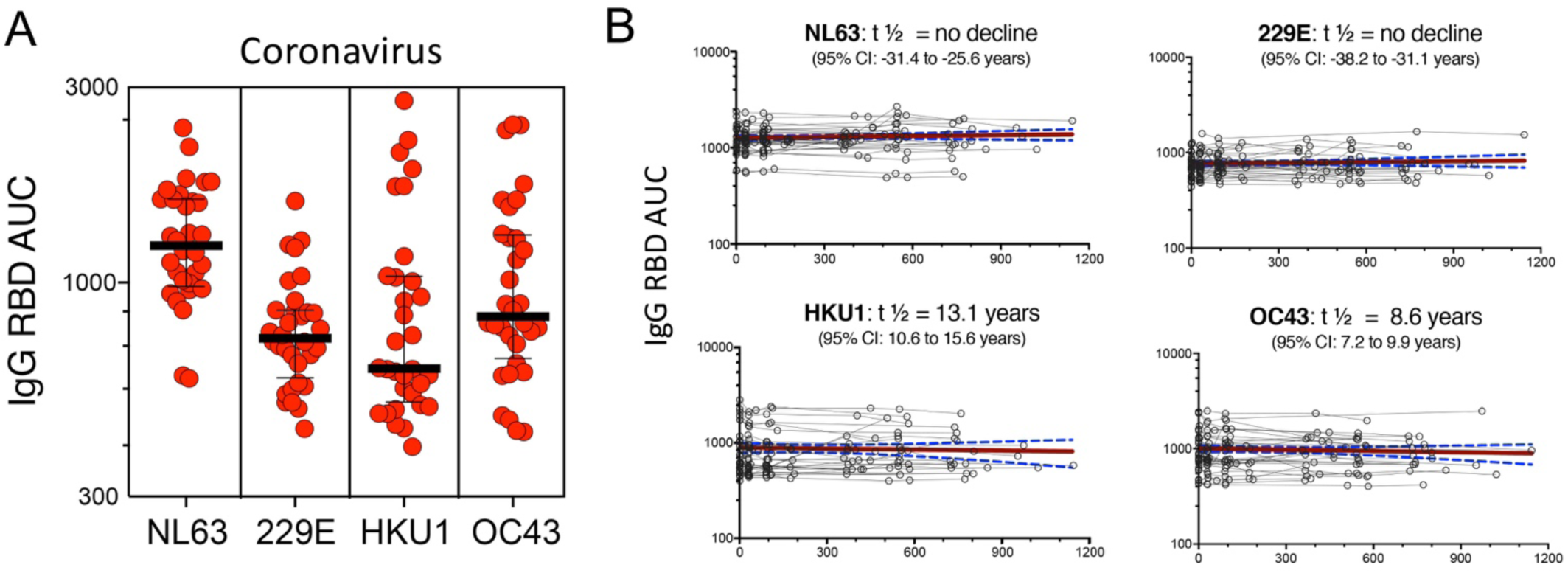
CCC-specific IgG responses are detected in all individuals and sustained overtime. (**A**) Plasma IgG titers, measured by the AUC, to the spike receptor binding domain (RBD) protein of the CCC viruses (HCoV-229E, HCoV-NL63, HCoV-HKU1 and HCoV-OC43) are shown for first time point of the longitudinal cohort (n=32). Geometric mean titers with SD are indicated. (**B**) Graphs show individual CCC antibody responses plotted for all time points of the longitudinal series and connected with lines for each subject (n = 32). The red line represents the median fitted curve from a nonlinear mixed effects model of longitudinal responses among those with a positive response at ≥ 1 time point, with 95%CI shown in blue dotted lines. t ½ calculated based on linear mixed effects model using R package nlme ^41^, t^1/2^ is shown as the median half-life estimated from the median slope with the associated 95% CI indicated.

### Correlation of CCC-specific CD4+ T cell and SARS-CoV-2 pre-existing responses

CCC have significant sequence homology to SARS-CoV-2. The cross-reactive CCC-specific T cell responses with SARS-CoV-2 ^26,27,46^ may contribute to modulation of SARS-CoV-2 infection and enhance responses to COVID-19 vaccination ^26,30–33^. We show that high CD4+ T cell memory OC43 reactivity is associated with higher levels of pre-existing memory reactivity to SARS-CoV-2 (**Fig. 6A)**. Similar patterns were observed for NL63, 229E and HKU1 (**Fig. S3)** but not for the unrelated and ubiquitous pathogen CMV (**Fig. 6B**). The highest levels of pre-existing SARS-CoV-2 reactivity were not associated with higher levels of HLA-DR/CD38+ expression (**Fig. 6C**) suggesting that SARS-CoV-2 cross-reactive cells are not the result of a recent activation or infection. Similarly, high pre-existing SARS-CoV-2 CD4+T cell responses did not correlate with higher IgG antibody reactivity (**Fig. 6D**). Overall, these results suggest that high CD4+ T cell reactivity to CCC, but not antibody reactivity, is predictive of cross-reactive SARS-CoV-2 CD4+ T cell responses.

**Fig. 6.**
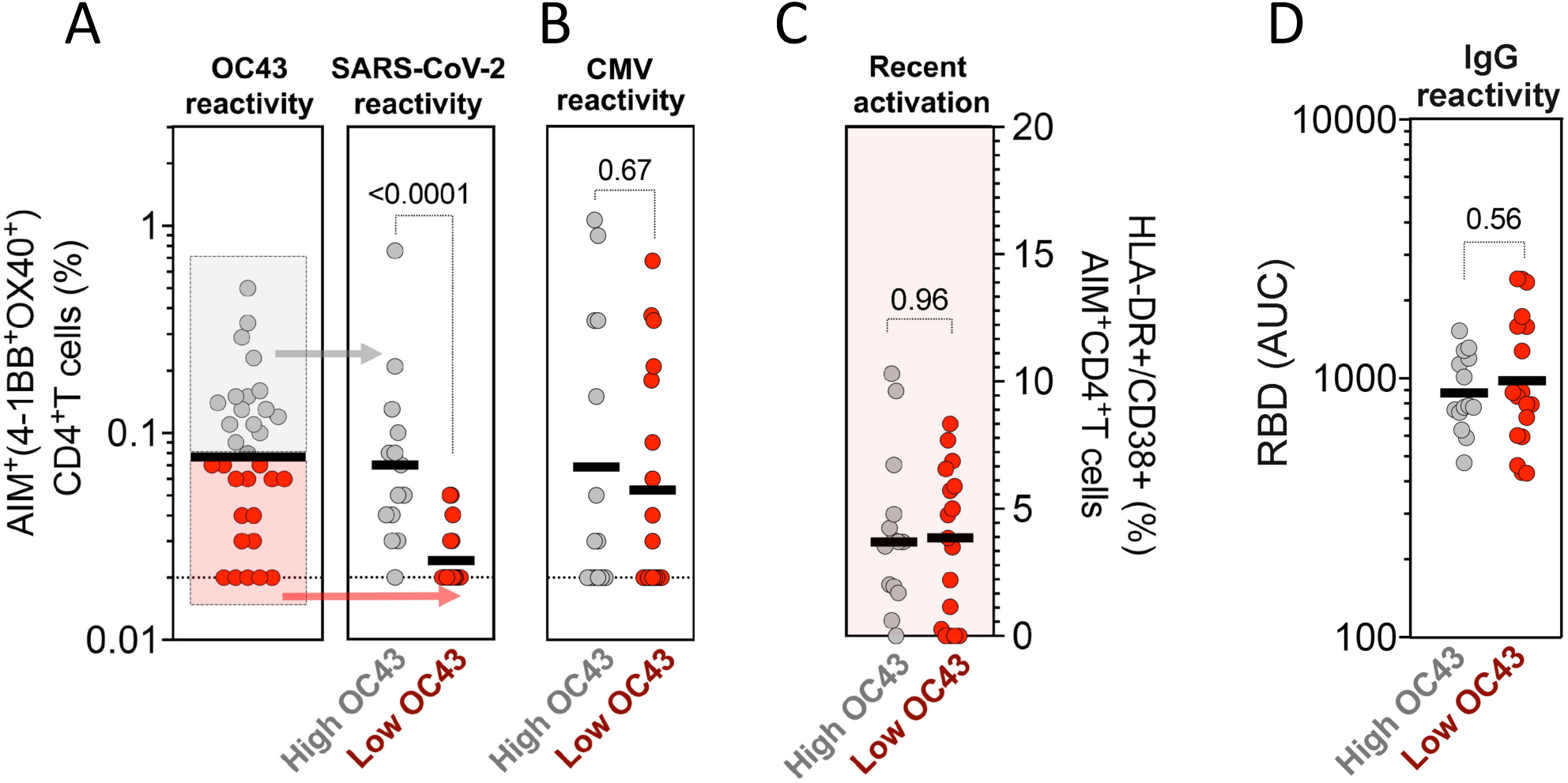
High CD4+ T cell reactivity to OC43 is associated with high pre-existing SARS-CoV-2 immunity. Antigen-specific T cell responses were measured as percentage of AIM+ (OX40+CD137+) CD4+ T cells after stimulation of PBMCs with peptides pools for (**A**) CCC (OC43) and SARS-CoV-2 (representing pre-existing immunity in pre-pandemic samples), (**B**) CMV as a control. (**C**) Recent activated CCC (OC43) specific T cell responses were measured by calculating the percent of HLA-DR^+^CD38^+^ of AIM+ (OX40^+^CD137^+^) CD4+ T cells. (**D**) Plasma IgG titers to CCC viruses (OC43) spike receptor binding domain (RBD) protein were measured by ELISA. (**A**-**D**) Each dot represents the response of an individual subject (n=32) at first time point with median bar shown. High responders for OC43 (above median bar in **A**) are shown in gray, and low responders for OC43 (below median bar in **A**) are shown in red. The different immune responses between high and low responders were compared using Mann-whitney test, and p values < 0.05 considered statistically significant.

## Discussion

Despite the high prevalence of CCC infection, most adults experience asymptomatic or mild common cold symptoms. Although seroconversion to CCC is near ubiquitous during childhood, little is known about the dynamics of CCC-specific memory responses in adults.

In this study, our goal was to characterize CD4+ T cell-specific memory responses to the four prototypic endemic and widely circulating CCC viruses in a longitudinal cohort, and using as a strategy ex vivo stimulation of PBMCs with peptides pools covering the entire proteome of each individual virus. In particular, our data shows that T cell reactivity against CCC is detected for most adult subjects, and that this reactivity is similar in magnitude to other antigens, sustained and relatively constant over time. These observations were paralleled by equally stable circulating Spike RBD antibody titers. While we cannot exclude that at least some reinfections might have occurred during the longitudinal follow up period, the preponderance of evidence suggest that the reactivity observed is associated with memory and persistent responses.

Previous work has suggested that substantial immune memory involving both arms of adaptive immune memory is generated against CCC ^17,23,25,26^. We observed that about 72-81% of subjects exhibited immune CD4+ T cell memory responses to each of the 4 CCC studied, consistent with frequencies detected in both community and heath care works cohorts ^26^, and similar to the findings of Tan et al. ^25^. Remarkably, we found a stable and sustained T cell and antibody response against CCC which are supported by recent experimental findings from Cohen et al. ^41^ or mathematical modeling ^19,20^ and argues against the short-lived nature of CCC responses ^16–18^. Our findings are also in agreement with the stability of responses against other viral infections, such as vaccinia or SARS-CoV-1 where antigen specific cells were detectable 50 years and 17 years post infection, respectively ^47,48^ and against TT which are remarkably stable for many years upon vaccination ^49^. It is quite possible that the CCC “steady state” reactivity might be result of both repeated infections in early childhood and occasional re-exposure and reinfection.

The data presented herein are also relevant in the context of CCC cross-reactivity with SARS-CoV-2. It has been shown that pre-existing T cell immunity elicited by past CCC exposures can influence COVID-19 responsiveness to vaccination and disease outcome ^30–33,35^. We found that while everybody had detectable antibody titers to CCC individuals varied in the level of T cell reactivity (possibly as a function of recent exposure, HLA type or other individual and environmental factors), and that the subjects with high CCC T cell reactivity, but not antibody titers, are those most likely to be associated with pre-existing SARS-CoV-2 immune reactivity. This is consistent with findings that CCC antibodies might not protect against SARS-CoV-2 infection or disease severity ^24,50^ but T cells do ^30,51^.

SARS-CoV-2 antibody and T cell effector activity contract over time and protection from infection wanes but protection from severe disease appears to be preserved in significant degree ^8,52–54^. Likewise, it has also been suggested that recurrent CCC infections are only rarely associated with moderate or severe clinical symptoms ^17,21^. In the context of the debate of the evolution of SARS-CoV-2 pandemic, our data are compatible with the notion upon enough people being exposed and vaccinated, SARS-CoV-2 at some point might turn endemic and resemble CCC, in terms of sustained herd immunity and relative protection not from infection but from symptomatic and severe disease. The development of pan-corona vaccines that target not only SARS-CoV-2 but also CCC viruses might contribute to further protection.

In summary, we found that in addition to widespread antibody reactivity to all the four CCC, memory T-cell responses are detected for most individuals and their reactivity remained stable and relatively constant over time. The characterization of the immune response to the prevalent and endemic CCCs provides a valuable reference for understanding the durability and eventual transition to an endemic state of SARS-CoV-2 in the aftermath of the pandemic.

### Limitations of the study

A limitation of this investigation is the unknown history of previous CCC exposure of the study participants. Assessment of CCC infection by RT-PCR was not part of the original study design for the study, and would also not have been logistically feasible over the longitudinal course of the study as it would have required frequent nasal swabs of all subjects. Therefore, in the present study, the stability of T cell and antibody responses could not be directly correlated with protection from symptomatic colds and/or infection. Furthermore, our analysis is limited to “steady state” responses in adults, and the evolution of CCC responses in children was not addressed. Additional limitations of this study are the relatively small cohort size investigated. Validation of the results in geographically distinct populations would be desirable to generalize the findings broadly.

## Acknowledgments

We wish to acknowledge all subjects for their participation and for donating their blood and time for this study. We are grateful to the La Jolla Institute for Immunology clinical core relentless efforts in obtaining blood samples. Research reported in this publication was supported by the National Institute of Allergy and Infectious Diseases (NIAID) of the National Institutes of Health (NIH) under Award numbers U01AI141995, U19AI142742 and U54CA260543, and contract number 75N93019C00065. The content is solely the responsibility of the authors and does not necessarily represent the official views of the National Institutes of Health.

## Author Contributions

Designing research studies, AS, RdSA, EDY; Investigation, AS, RdSA, EDY, TN, EW, EG, LP, JM; Data Analysis, EDY, RdSA, LP; Resources, AS and RdSA; Manuscript writing, AS, RdSA, EDY, LP, DW, AG; Supervision, AS and RdSA; Project Administration, AF; Funding Acquisition, AS and RdSA.

## Materials and methods

### Study cohort and PBMC isolation

The purpose of this study was to investigate the immunological memory to common cold corona viruses in a longitudinal cohort collected during pre-pandemic time. Blood donations from 32 donors previously recruited in a *Bordetella Pertussis* observational study were collected under IRB approved protocols at the La Jolla Institute for Immunology (protocol no. VD-101), before COVID-19 pandemic from 2016-2019. There were 3 to 7 longitudinal blood donations per donor spanning time periods from 6 months to more than 3 years. Each participant provided informed consent and was assigned a study identification number with clinical information recorded. Exclusion criteria included pregnancy at the start of the study; presentation of severe disease; medical treatment that might interfere with study results and/or antibiotic use or fever (>100.4°F [38°C]). No exclusions were made due to race, ethnicity, or gender. In all cases, PBMCs were isolated from whole blood by density gradient centrifugation according to manufacturer instructions (Ficoll-Hypaque, Amersham Biosciences, Uppsala, Sweden) and cryopreserved for further analysis.

### Synthesize of epitope pools

In the current study, in order to investigate the CCC and SARS-CoV-2 specific T cell responses, we used megapools (MPs) combing the overlapping spike (S) epitope pools and predicted HLA class II CD4+ T cell epitope pools from the rest of the genome (R) (**Table 2**), generated using previously described strategies ^26,36^. We also studied antigen-specific responses against a panel of other respiratory viruses (influenza, RSV, and rhinovirus), chronically infectious viruses (EBV, CMV, and VZV), and ubiquitous bacterial vaccine antigens (TT and PT) using peptide sets described in **Table 2**. Detailed overall information of the MPs composition, peptide numbers as well as references are specified in **Table 2**. Individual peptides were synthesized by TC peptide lab (San Diego, CA) and pooled by protein combinations and resuspended to a final concentration of 1 mg/mL in DMSO.

### CCC Spike protein RBD Enzyme-Linked Immunosorbent Assay

All plasma samples tested by ELISA assay were heat-inactivated at 56°C for 30 min to reduce risk from any possible residual virus in serum. Briefly, 50 µL of Streptavidin (Invitrogen) at 4 µg/mL in Tris-Buffered Saline (TBS) pH 7.4 was coated in the 96-well, high-binding microtiter assay plate (Greiner Bio-One cat # 655061) for 1 hour at 37°C. The coating solution was removed, then 100 µL of blocking solution, 1:1 Non Animal Protein-BLOCKER™ (G-Biosciences) in TBS was added for 1 hour at 37°C. Serum samples were serially diluted (1:40 – 1:8100), in 3% Bovine Serum Albumin (BSA) in TBS containing 0.05% Tween 20 (TBST) with respective biotinylated spike RBD antigens from CCC at 1 µg/mL in a 96-round-well V bottom plate (Diaago cat # R96-300V) and incubated for 1 hour at 37°C. The blocking solution was removed, then 50 µL of diluted serum was added to the assay plate and incubated for 15 minutes at 37°C. The plate was washed three times using wash buffer (1X TBS containing 0.2% Tween 20), then 50 µL of horseradish peroxidase-conjugated secondary Goat Anti-Human secondary IgG antibody (Cat No: 109-035-008, Jackson ImmunoResearch) at 1:40,000 dilution in 3% milk was added for 1 hour at 37°C. The plate was washed three times using wash buffer, then 50 µL of 3,3’,5,5’-Tetramethylbenzidine (TMB) Liquid Substrate (Sigma-Aldrich) was added to the plate, and absorbance was measured at 450 nm using a plate reader (Molecular Devices SpectraMax ABS Plus Absorbance ELISA Mcroplate Reader) after stopping the reaction with 50 µl of 1 N HCl. Area under the curve for titration experiments for each sample were calculated by the trapezoidal model implemented in Prism Version 9.3.0.

### Activation induced cell marker (AIM) assay

The AIM assay was performed as previously described ^27^. Cryopreserved PBMCs were thawed by diluting the cells in 10 mL complete RPMI 1640 with 5% human AB serum (Gemini Bioproducts) in the presence of benzonase [20 ml/10ml]. Cells were cultured for 20 to 24 hours in the presence of CCC or SARS-CoV-2 specific and other common antigen pools (1ug/ml) in 96-wells U bottom plates with 1×10^6^ PBMC per well. An equimolar amount of DMSO was added as a negative control and phytohemagglutinin (PHA, Roche (San Diego, CA) 1 mg/ml) was used as the positive control. Cells were stained and activation of CD4+ T cell measured by the CD137 and OX40 marker combination. The detailed information of all the antibodies used are summarized in **Table S1**. All samples were acquired on a ZE5 cell analyzer (Biorad laboratories, Hercules, CA) and analyzed with FlowJo software (Tree Star, Ashland, OR). AIM+ CD4+ T cells data were calculated as percent of total CD4+ T cells background subtracted or stimulation index. Background subtracted data were derived by subtracting the percentage of AIM+ cells percentage after each MP stimulation from the DMSO stimulation. The Stimulation Index (SI) was calculated by dividing the count of AIM+ cells after SARS-CoV-2 pools stimulation with the ones in the negative control. A positive response was defined as SI greater than 2 and AIM+ response above the threshold of positivity after background subtraction. The limit of detection (0.02%) was calculated based on 2 times 95% CI of geomean of negative control (DMSO), and the threshold of positivity (0.03%) was calculated based on 2 times standard deviation of background signals according to previous published studies ^27,52^. All data below 0.02 or SI<2 were set to 0.02 or 2 for plotting and statistical analysis. The detailed gating strategy used to define CD4+ AIM reactive cells (OX40^+^CD137^+^), memory (CD45RA/CCR7), and activated sub-populations (HLA-DR+CD38+) is listed in **Fig. S4**. Gates were drawn relative to the unstimulated condition for each donor.

### Statistical analysis

Experimental data were analyzed by GraphPad Prism Version 9 (La Jolla, CA) and Microsoft Excel Version 16.16.27 (Microsoft, Redmond, WA). The statistical details of the experiments are provided in the respective figure legends. Data were analyzed by Mann-Whitney test (two-tailed) to compare between two groups, and Kruskal-Wallis test adjusted with Dunn’s test for multiple comparisons to compare between multiple groups. The regression lines and estimated t ½ was calculated based on linear mixed effects model using R package nlme as previously described by Cohen et al. 2021 ^41^. Data were plotted as geometric mean with geometric SD for log scale and median with interquartile range for numeric scale. *p* values < 0.05 (after adjustment if indicated) were considered statistically significant.

### Study approval

This study was approved under IRB protocol approval (VD-101) at the La Jolla Institute for Immunology. All donors were able to provide informed consent, or had a legal guardian or representative able to do so. Each participant provided informed consent and was assigned a study identification number with clinical information recorded.

## Materials & Correspondence

Epitope pools used in this study will be made available to the scientific community upon request, and following execution of a material transfer agreement (MTA), by contacting A.S. and R.d.S.A. Likewise, biomaterials archived from this study may be shared for further research with MTA.

## Data and code availability

The datasets generated and analyzed in this study will be shared by the corresponding authors upon reasonable request. This paper does not report original code.

