## Supplementary Figures and Tables for "Immunological memory to Common Cold Coronaviruses assessed longitudinally over a three-year period"

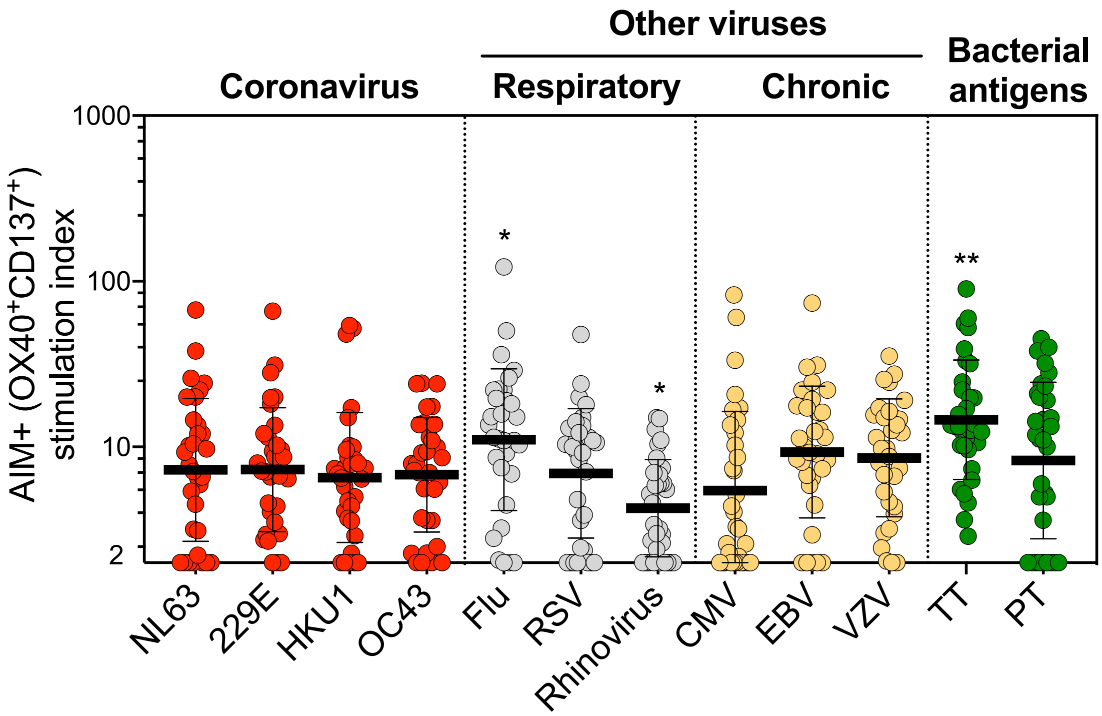


**Fig. S1 Stimulation Index of CD4+ T cell responses to four representative CCC and other pathogens.** Common cold coronavirus (CCC) and several other human pathogens-specific T cell responses were measured as percentage of AIM+ (OX40+CD137+) CD4+ T cells after stimulation of PBMCs with peptides pools. Graphs show individual response of four CCC (NL63, 229E, HKU1, and OC43) and other pathogens plotted as stimulation index (SI) against DMSO negative control. First time point of the longitudinal series is plotted (n = 32). Data are represented as geometric mean and SD. Kruskal-Wallis test adjusted with Dunn's test for multiple comparisons was performed and adjusted p values < 0.05 considered statistically significant. *, $p < 0.05$, **, $p < 0.01$.

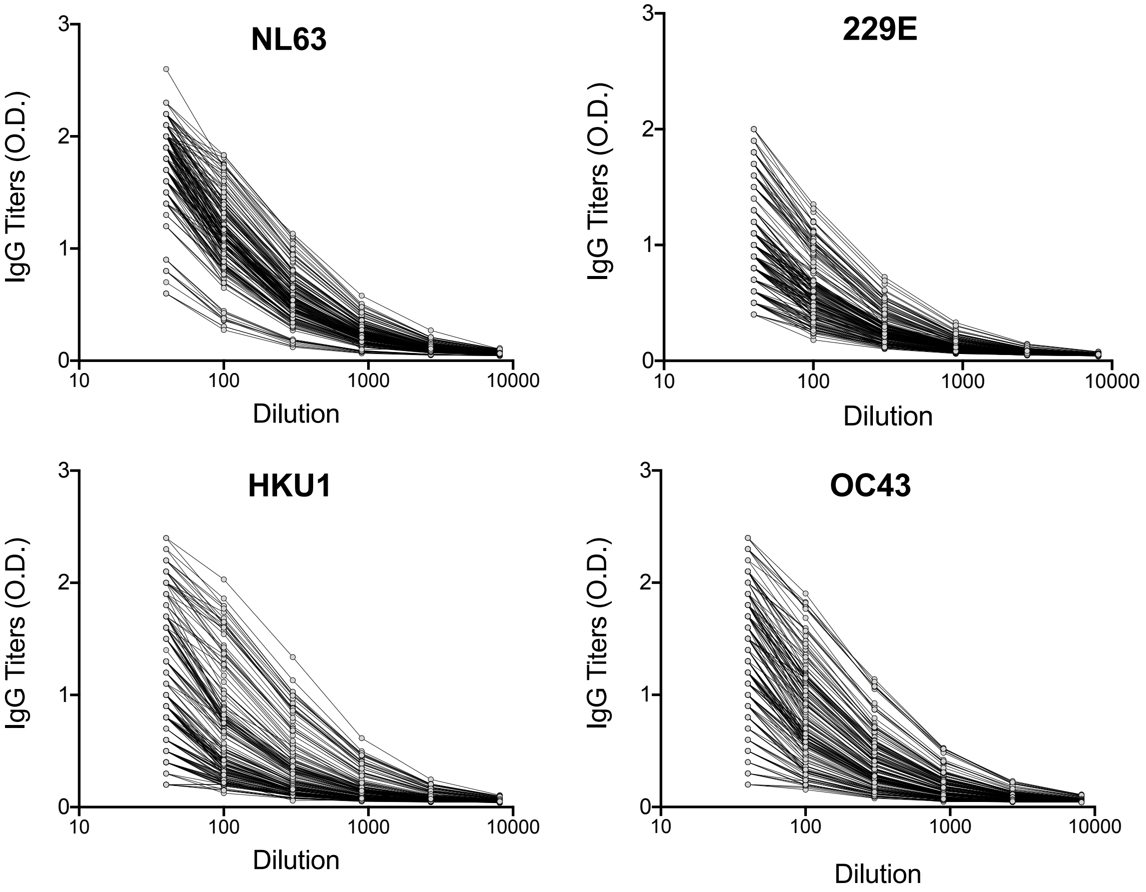


**Fig. S2 IgG serial dilutions for endpoint titers and AUC calculation.** Plasma ELISA IgG serial dilutions to calculate the area under the curve (AUC) for CCC viruses (229E, NL63, HKU,1 and OC43) spike receptor binding domain (RBD) protein are shown for the longitudinal cohort (n = 32).

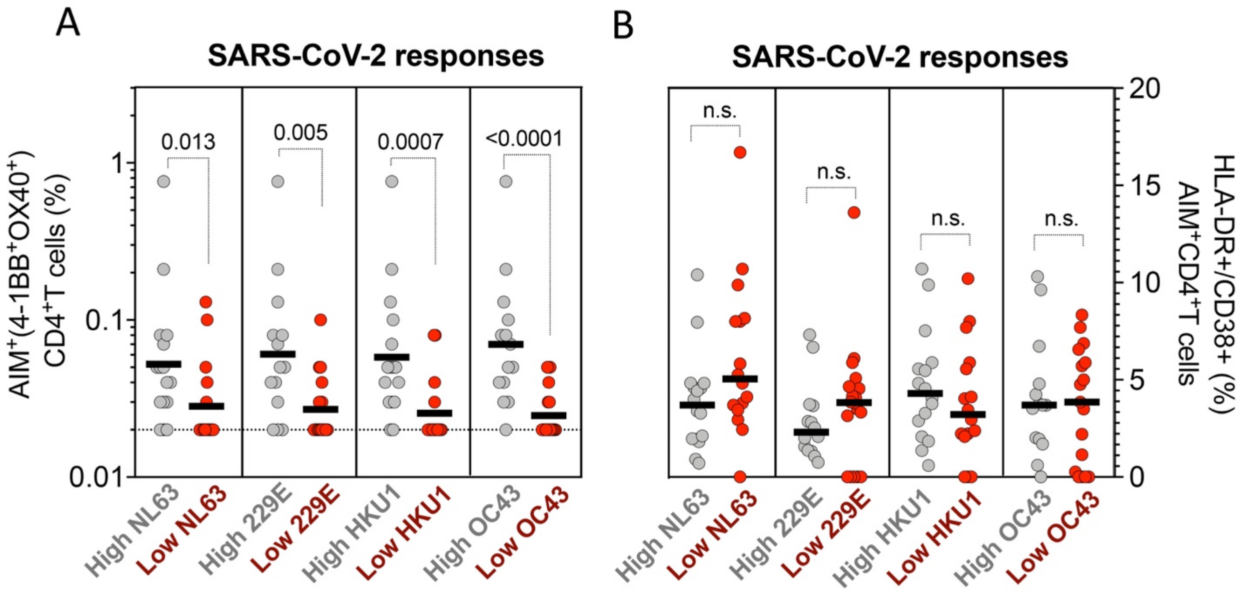


**Fig. S3 High CD4+ T cell reactivity across all the CCC viruses is associated with high pre-existing SARS-CoV-2 immunity.** (**A**) Antigen-specific T cell responses were measured as percentage of AIM+ (OX40+CD137+) CD4+ T cells after stimulation of PBMCs with peptides pools for CCC and SARS-CoV-2 (representing pre-existing immunity in pre-pandemic samples). (**B**) Recent activated CCC-specific T cell responses were measured by calculating the percent of HLA-DR$^{+}$CD38$^{+}$ of AIM+ (OX40$^{+}$CD137$^{+}$) CD4+ T cells. Each dot represents the response of an individual subject (n=32) at first time point. Median is shown. High responders for each CCC are shown in gray, and low responders in red. The different SARS-CoV-2 specific immune responses between high and low CCC responders were compared using Mann-whitney test, and p values < 0.05 considered statistically significant.

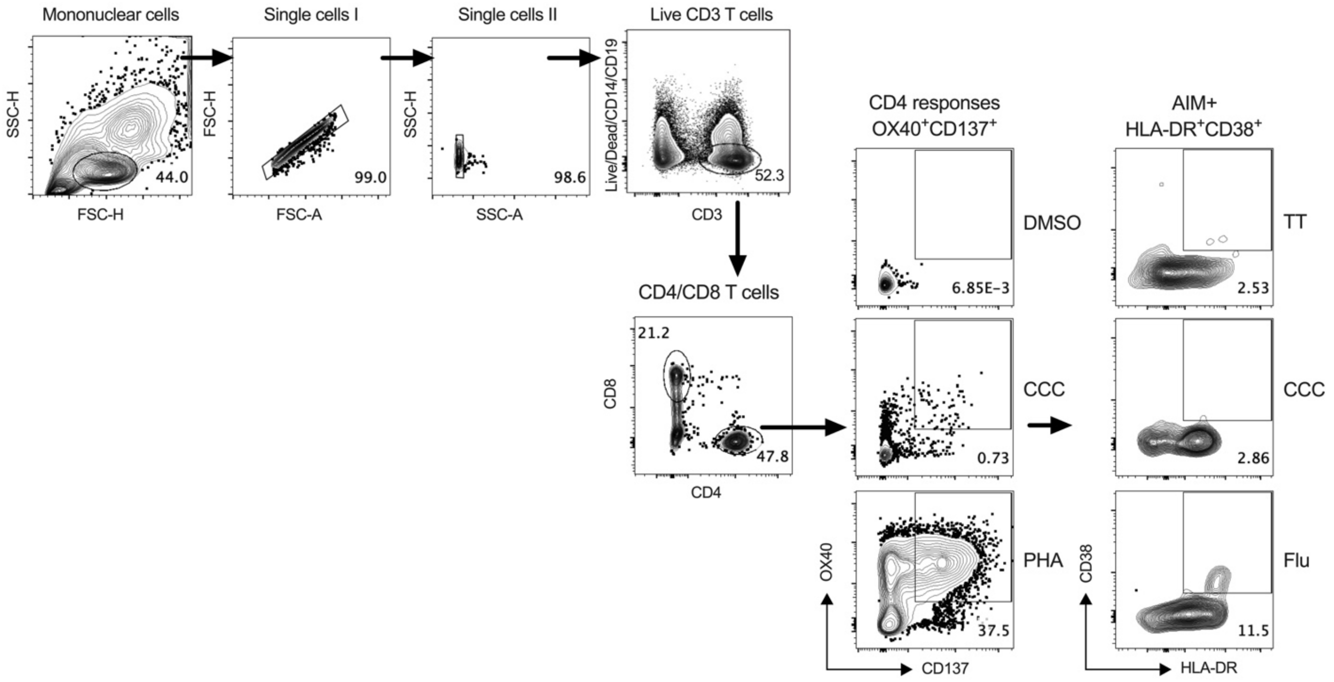


**Fig. S4 Ilustrative flow cytometry gating strategy for the assessement of antigen-specific CD4+ T cell responses.** Representative gating of reactive OX40+CD137+ CD4+ T cells (AIM+) from donor PBMCs is shown. Briefly, mononuclear cells were gated out of all events followed by subsequent singlet gating. Live CD3+ cells were gated as Live/Dead-CD14-CD19-CD3+. Cells were then gated as CD4+CD8-, and antigen-specific cells definied as OX40+CD137+ CD4+ T cells (AIM+) and after antigen stimulation, and frequencies calculated as percent of total CD4+ T cells. Representative AIM+ responses after stimulating with positive (PHA) or negative (DMSO) controls and CCC (OC43) specific megapools are presented on the right. Recently activated cells (HLA-DR+CD38+) were further gated from AIM+ cells. Representative plots are show for TT, CCC (OC43) and Flu.

**Table S1.** List of the antibodies used in this study

| Antibody | Fluorochrome | Clone | Vendor | Catalog number |
|---|---|---|---|---|
| CD3 | BV805 | UCHT1 | BD Biosciences | 612895 |
| CD4 | BV605 | RPA-T4 | BD Biosciences | 562658 |
| CD8 | BV496 | RPA-T8 | BD Biosciences | 612942 |
| CD14 | V500 | M5E2 | BD Biosciences | 561391 |
| CD19 | V500 | HIB19 | BD Biosciences | 561121 |
| CD137 | APC | 4B4-1 | Biolegend | 309810 |
| OX40 | PE-Cy7 | Ber-ACT35 | Biolegend | 350012 |
| CD69 | PE | FN50 | BD Biosciences | 555531 |
| HLA-DR | AF700 | LN3 | eBiosciences | 56-9956-42 |
| CD38 | BV786 | HIT2 | BD Biosciences | 563964 |
| CD45RA | BV421 | HI100 | Biolegend | 304130 |
| CCR7 | FITC | G043H7 | Biolegend | 353216 |
| Live/Dead Viability | eF506/Aqua | - | eBiosciences | 65-0866-18 |
